# The association between adiposity and inpatient hospital costs in the UK Biobank cohort

**DOI:** 10.1101/399600

**Authors:** Padraig Dixon, George Davey Smith, William Hollingworth

## Abstract

**Background:** High adiposity is associated with higher risks for a variety of adverse health outcomes, including higher rates of age-adjusted mortality and increased morbidity. This has important implications for the management of healthcare systems, since the endocrinal, cardiometabolic and other changes associated with increased adiposity may be associated with substantial healthcare costs.

**Methods:** We studied the association between various measures of adiposity and inpatient hospital costs through record linkage between UK Biobank and records of inpatient care in England and Wales. UK Biobank is a large prospective cohort study that aimed to recruit men and women aged between 40 and 69 from 2006 to 2010. We applied generalised linear models to cost per person year to estimate the marginal effect and averaged adjusted predicted cost of adiposity on inpatient costs.

**Results:** Valid cost and body mass index (BMI) data from 457,689 participants were available for inferential analysis. Some 54.4% of individuals included in the analysis sample had positive inpatient healthcare costs over the period of follow-up. Median hospital costs per person year of follow-up were £89, compared to mean costs of £481. Mean BMI overall was 27.4 kg/m2 (standard deviation 4.8). The marginal effect of a unit increase in BMI was £13.61 (99% confidence interval: £12.60 to £14.63) per person year of follow up. The marginal effect of a standard deviation increase in BMI was £69.20 (99% confidence interval: £64.98 to £73.42). The marginal effect of becoming obese was £136.35 (99% confidence interval: £124.62 to £148.08). Average adjusted predicted inpatient hospital costs increased almost linearly when modelled using continuous measure of adiposity. Sensitivity analysis of different scenarios did not substantially change these conclusions, although there was some evidence of attenuation of the effects of adiposity when controlling for waist-hip ratios, and when individuals who self-reported any pre-existing conditions were excluded from analysis.

**Conclusions:** Higher adiposity is associated with higher inpatient hospital costs. Further scrutiny using causal inferential methods is warranted to establish if further public health investments are required to manage the large healthcare costs observationally associated with overweight and obesity.

## INTRODUCTION

Body mass index (BMI) – weight divided by the square of standing height – is a widely used indicator of nutritional status, adiposity and overall health (1-7). Longitudinal, cross-country data reveal both a recent increase in mean BMI levels for men and women, and increased variance in BMI (8). Globally, more individuals are becoming obese (BMI>=30kg/m2) than are transitioning out of underweight status (BMI<=18.5kg/m2) (8, 9), leading to marked increases in the absolute numbers of individuals in the former category. Some 2.1 billion individuals throughout the world were overweight (BMI>25kg/m2) or obese in 2013, an increase of 1.4 billion from 1980 (4). This reflects a global prevalence of overweight or obesity amongst men of 28.8% and 29.8% amongst women (4).

The underlying aetiology between BMI and health is complex (10), but robust associations have nevertheless been found between higher BMI and risks for a variety of adverse health and social outcomes, including higher rates of age-adjusted mortality (11), increased morbidity (8, 12) and poor labour market outcomes (13). This has important implications for the management of healthcare systems (14), since the endocrinal (15), cardiometabolic (16, 17) and other changes (14) associated with increased adiposity are associated with substantial healthcare resource requirements (18).

Cawley et al (19) estimate that the direct medical care costs of obesity of US adults amounted to 27.5% of health expenditures for non-institutionalized US adults, and noted adult obesity has become more expensive over time in the US due to both increased prevalence and higher treatment costs. Kent et al (20) attribute 14.6% of total annual hospital costs in women aged 55-79 in England to overweight and obesity, with this figure rising to 52% for individuals with BMI >40kg/m2.

This article assesses the observational association between BMI and other measures of adiposity with admitted patient hospital costs using individual-level data from UK Biobank, a large prospective cohort study. This is the first study to develop cost estimates for UK Biobank, the newly created data for which will be returned to the cohort and freely available for other researchers to use. Observational analyses of the type deployed in this paper cannot answer causal questions, or identify the mechanisms that mediate the association between adiposity and healthcare costs, but do have utility in documenting policy-relevant associations, offering context for previous observational and causal studies, and providing evidence for meta-analysis.

A further advantage of this analysis is that the results may be triangulated (21) with other sources of evidence with potentially orthogonal sources of bias to come to an overall view about the association between BMI and healthcare costs. Together, diverse sources of evidence, of which the present analysis is one source, can be used to inform the cost-effectiveness of interventions and policies targeting the prevention and treatment of obesity. As Wang et al (14) note, “A systematic understanding of the potential morbidity and cost implications of specified hypothetical changes in body-mass index trajectories…is crucial for formation of effective and cost-effective strategies, establishment of research and funding priorities, and creation of the political will to address the obesity epidemic.”

## METHODS

### UK Biobank – features of study design and participants

Adults aged between 40-69 and living within approximately 25 miles (40km) of 22 assessment centres in England, Wales and Scotland were invited to participate in the UK Biobank study. Some 9.24 million individuals registered with the UK’s National Health Service (NHS) were sent postal invitations, and 503,317 individuals ultimately joined the study cohort (22), for a response rate of approximately 5.45%. Participation rates were higher for women (6.4%) than men (5.1%) (22). The cohort is not representative of the national population from which it is drawn, being wealthier, healthier and better educated. The implications of the response rate and the representativeness of the cohort are considered in the Discussion section below. Ethical approval for UK Biobank was received from the North West - Haydock Research Ethics Committee (reference: 11/NW/0382).

Participant assessments, undertaken at the 22 centres between 2006 and 2010, comprised consent, self-completion of an electronic questionnaire, computer-assisted interviews, specimen collection and measurement of physical function (23). Individual participant baseline data was linked with participant consent to, inter alia, death registers and records of certain forms of care in NHS hospitals.

### Measurement of body mass, waist-hip ratios and bio-impedance

Three different but cognate measures that indicate adiposity were measured at the baseline appointment: BMI measured using height and weight, bio-impedance (opposition in biological tissues to alternating current) and circumferences of the waist and hip.

Weight was measured following the removal of shoes and heavy outer clothing using Tanita BC-148MA body composition analysers. Participants’ standing shoeless height was measured using a Seca 202 device. These measurements were used to create an index of body mass in terms of kg/m2.

The built-in algorithms of the Tanita analysers were also used to estimate body composition following measurement of bio-impedance. Mass measured using impedance was calculated in increments of 0.1kg. A separate measure of adiposity was calculated from these impedance measures by dividing by standing height of participants to create another index of body mass, also in terms of kg/m2.

These two measures of adiposity were highly concordant (Lin’s rho=0.99996, p<0.00001) with a mean difference between traditional and impedance measures of −0.01 (99% confidence interval: −0.12 to 0.10). Impedance based BMI was therefore used where the traditional BMI measure was missing. A total of 685 observations had a mean difference between traditional and impedance-based measures of BMI of more than 5 standard deviations from the mean difference and were excluded from the analysis.

Waist circumference (at the umbilicus) and hip circumference were measured in centimetres using a Wessex sprung tape measure. A waist-hip ratio (WHR) was calculated from these measurements to reflect central adiposity (24, 25). We estimated separate models including WHR as a continuous variable and as a binary variable. The binary variable was constructed to reflect measures of regional adiposity using a ratio>0.85 for women and >0.9 for men. It is important to note that while these binary cut-off points are used in a number of countries, there is no widely accepted definition of these cut-offs, despite the frequent attribution of these figures to World Health Organization reports (see and particularly the discussion in (26)). Nevertheless, this classification has utility as a comparison to the binary cut-off of obesity defined for the conventional BMI measures, as well to published literature using these or similar classifications for WHR measures.

As a sensitivity analysis, we estimated models of the continuous WHR outcome using continuous BMI as a covariate, and vice versa. These models estimated, respectively, BMI-adjusted WHR, and WHR-adjusted BMI outcomes conditional on all baseline covariates, and each model assumes that WHR is not on the causal pathway between BMI and hospital costs and vice versa.

### Measurement of costs

Admitted patient care episodes, sometimes referred to as inpatient care episodes, were obtained from linked Hospital Episode Statistics (HES) for English care providers and from Patient Episode Database for Wales (PEDW) for Welsh providers. These two data sources are very similar in structure and purpose, and were analysed as a single dataset following reconciliation where required. Inpatients are those who are admitted to hospital and occupy a hospital bed but need not necessarily stay overnight (i.e. day case care).

Episodes of care refer to a period of care in which a patient is under the care of a single consultant working at a single hospital provider. An admission may comprise just one “Finished Consultant Episode” (FCE), or many such episodes if the patient receives care from more than one consultant.

Each FCE is associated with information on the patient, consultant, and hospital overseeing their care. For example, FCEs are associated with procedures undertaken (based on OPCS-4 procedure codes and diagnoses made (based on ICD-10 diagnosis codes (27)). These FCEs were converted into Healthcare Resource Groups (HRGs) by using data on patient characteristics, length of stay, procedures, diagnoses, and other information by using NHS “Grouper” software (28).

HRGs reflect groups of similar patient activity, and are used in England and Wales as the basis for casemix-adjusted remuneration of publicly funded NHS hospitals. In turn, these HRGs were cross-classified against NHS Reference unit costs, allowing costs to be assigned to each FCE. These costs were calculated to reflect differences between categories of care included in inpatient care, such as elective and non-elective episodes, and accounted for the additional costs associated with patients undergoing long hospital stays. Costs were separately calculated for so-called “unbundled” elements of care, such as diagnostic imaging, for which elements of cost and care activity are represented separately from other elements of inpatient care.

Due to differences in the collection and valuation of care in Scottish hospitals compared to hospitals in England and Wales, only costs from the latter two jurisdictions are included in this analysis. Costs were calculated for episodes of care occurring on or after 1 April 2006 to reflect major changes to the hospital payment system that came into effect on that date (29). Participants who attended UK Biobank baseline appointments before this date were removed from the analysis.

Remaining cohort members therefore report person years from recruitment, and contribute data to HES until death, emigration, or 31 March 2015, the current censoring date for linked HES data. Note that the event of death does not constitute censoring in this analysis, since the number of all care episodes and associated costs is known following death. Emigration out of the UK amongst the cohort is reported to be a modest 0.3% (22) and is not accounted for in our analysis. We do not have information on patients who may have received inpatient care in Scottish hospitals while living in England or Wales. Costs were converted into a per patient cost by summing across all episode costs per person-year of follow-up in constant 2016/17 prices.

### Covariates

All models adjusted for covariates obtained from participant responses at the baseline appointment at UK Biobank assessment centres. These covariates were sex, age at recruitment, days per week of vigorous physical activity (categorical ranging from 0-7), frequency of alcohol intake (categorical ranging from “Daily or almost daily” to “Never”), educational and professional qualifications (categorical), employment status (categorical, ranging from “In paid employment or self-employed” to “Full or part-time student”), and a measure of deprivation calculated using the Townsend score converted into quintiles.

To assess the effects of residual confounding and of reverse causality from pre-diagnostic or prodromal disease (7), we conducted sensitivity analysis restricting the analysis to include only “never smokers” (n=249,423) and including only individuals reporting no pre-existing illness at baseline (n=106,388). Smoking status is self-reported and may exhibit measurement error as a consequence (30). It is possible that focusing only on never smokers (who are probably less likely to mis-report their smoking status than current or former status) may avoid some of the confounding that may otherwise be present in the BMI-hospital cost association. However, the causal relationship between BMI, smoking and healthcare costs is complicated and very likely bidirectional (31)(32). It is important to emphasise that accounting for residual confounding and reverse causality in this way is necessarily incomplete given the observational study design.

Including only those individuals with no health conditions at baseline may partially capture the impact of reverse causality if individuals with many pre-existing conditions are on treatment regimens that raise or lower BMI, so that the direction of causality runs from costs to BMI. Again, this is a partial attempt at accounting for reverse causality and reflects only baseline information that was accurately self-reported.

Two covariates (the exercise variable and the qualifications variable) had slightly higher rates of missing data than others, and we conducted a post hoc sensitivity analysis excluding these two variables from the adjusted regression models.

### Statistical methods

The primary objective of our analysis was to predict total admitted patient hospital costs as a function of the conditional marginal effect of body mass on healthcare costs. Marginal effects for categorical and binary adiposity-related variables refer to the effects of a change in category, e.g. from “overweight” to “obese”. Marginal effects for continuous outcomes were calculated for specific changes – unit changes and standard deviation changes – in the adiposity related variable. The magnitude of these effects depends on the values of other covariates included in the model – these values of other covariates were left at their observed values to calculate an average marginal effect.

The average adjusted predictions that follow from this calculation adjust for variation in these other covariates. These predicted costs therefore do not necessarily reflect the hypothetical costs of an “average” individual, nor necessarily the expected costs for any specific individual in the sample. Average adjusted predicted costs were calculated for the entire analysis sample, for the samples defined in sensitivity analyses, and at representative ages stratified by sex.

Cost is always non-negative, the modal value is often zero, and the distribution of healthcare costs tends to be skewed with a long right tail reflecting very high expenditures incurred by a small number of individuals. Linear models may predict negative costs for some individuals, may not estimate mean effects without bias in the presence of long tails, and are likely to be inefficient in the presence of heteroskedasticity. Logarithmic transformations can address skewness to some extent but are not well suited for handling zero cost observations, and require transformations of regression output back to the original scale, a process which itself may cause bias in the presence of heteroskedasticity (33).

Generalised linear models (GLM) were therefore used to estimated average adjusted predicted costs: rather than modelling the conditional expectation of the logarithm of cost, the logarithm of the conditional expectation of cost was estimated, and the relationship of the mean to variance in the outcome data was assessed and modelled. A so called “single-index model” was used, in which the zero-cost and positive-cost outcomes were modelled with a single density function. As Deb (34) et al note, this approach is appropriate when the object of inferential interest is related to the mean function of the outcome, conditional on covariates, such as the marginal effect of a covariate on mean outcomes. This type of inference is the focus of this paper, which is concerned with the marginal effects of measures of adiposity on inpatient hospital costs. However, in sensitivity analysis, we also estimated models under a two-part model which decomposed the density of the outcome into a mixture of two parts: one that models the zero cost observations (estimated with a logit model), and one that models the non-zero cost observations using a GLM structure.

Two tests, neither necessarily definitive, were used to identify an appropriate link function for all models. Box-Cox tests on positive cost observations were used to identify scalar powers that resulted in the most symmetric transformed distribution. A Pregibon-type (35) link test was performed to check whether including the squared outcome variable as a covariate had explanatory value above models excluding this term. To inform the choice of the family function, a modified version of the Park test was used to characterise the relationship between the variance of the error term and the expected value of the cost outcome variable. All models use robust standard errors to allow for potential mis-specification of the link and family functions.

All analyses were conducted in Stata version 15.1 (StataCorp, College Station, Texas). Code used in the analysis is available at github.com/pdixon-econ.

## RESULTS

Following exclusion of data as described above, records from up to 457,689 participants were available for the inferential analysis (Figure 1), of whom 54.4% at baseline were female (Table 1). Person years of follow-up ranged from 4.5 to 9 years (mean and median=6.1 years). Mean age at baseline for women was 56.3 years, and 56.7 years for men. Median hospital costs per person year of follow-up were £89, compared to mean costs of £481, reflecting the highly skewed distribution that is charactristic of this type of cost data.

**Figure 1.**
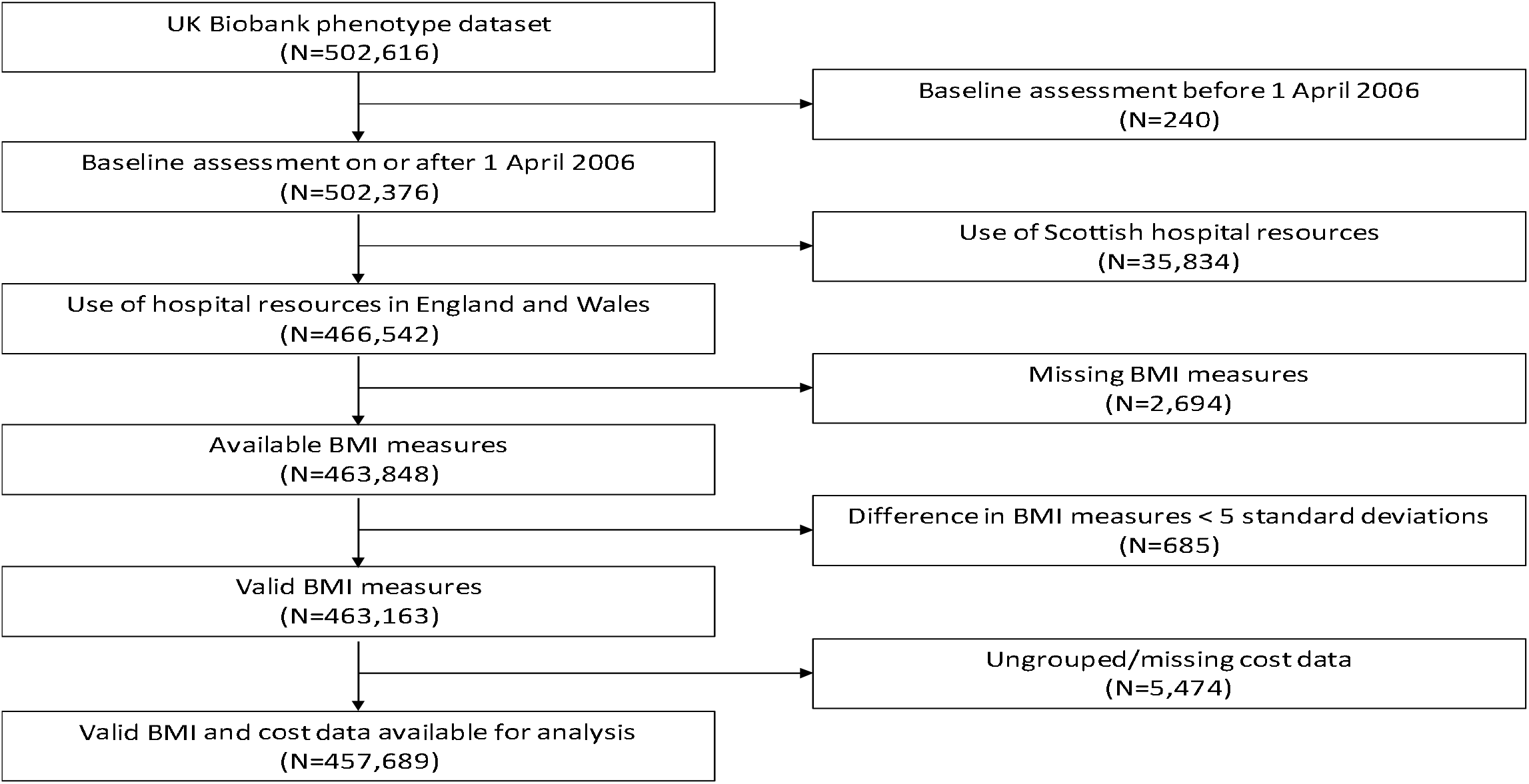
Flow chart of participants available for analysis. Notes to Figure: BMI: Ungrouped cost data refers to data for which HES data is not sufficient to assign a Healthcare Resource Group. This typically where the primary diagnosis was not specific and was coded to an unspecified injury (or similar) classification.

**Table 1.**
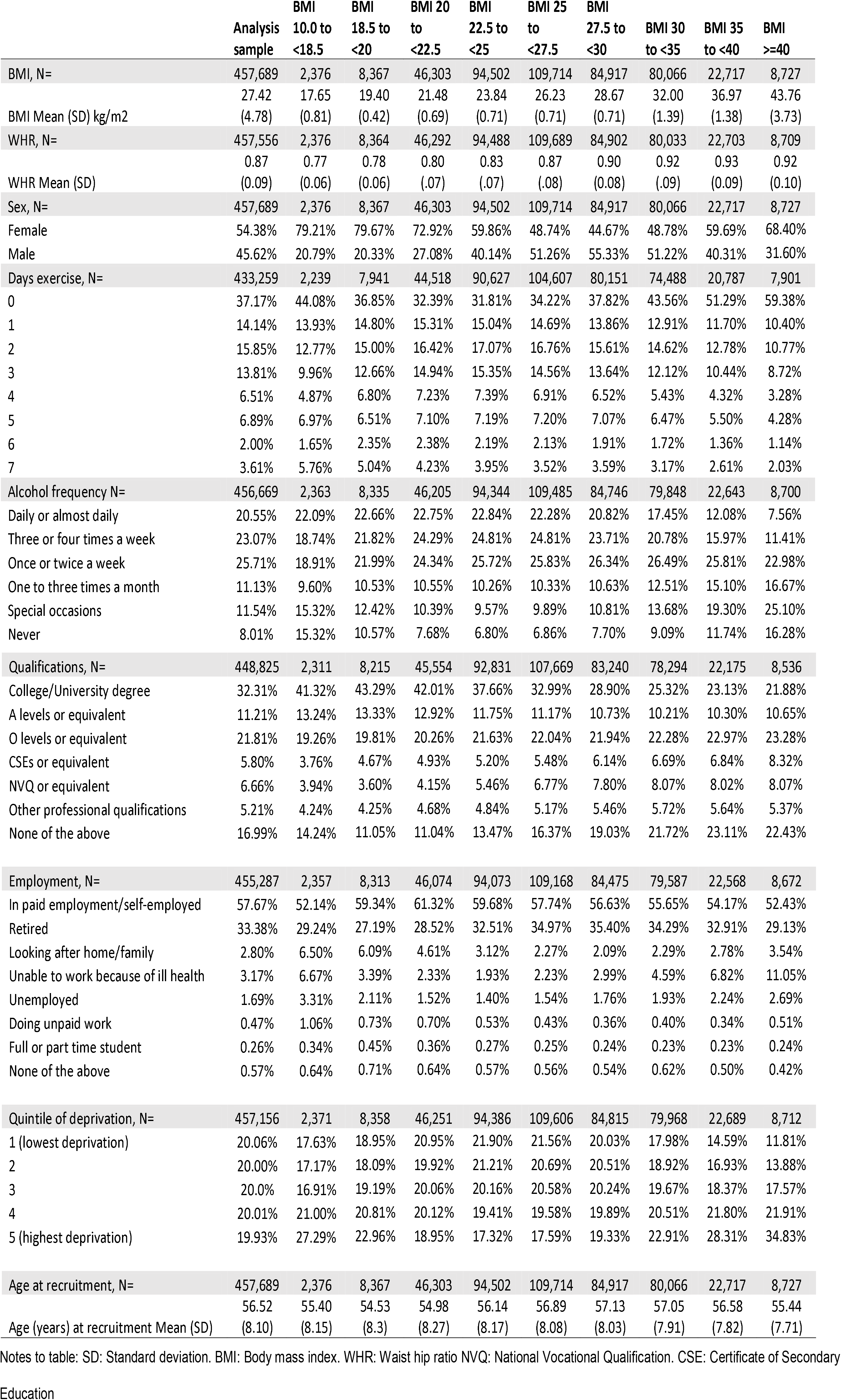
Descriptive statistics for analysis sample.

|  | Analysis sample | BMI 10.0 to <18.5 | BMI 18.5 to <20 | BMI 20 to <22.5 | BMI 22.5 to <25 | BMI 25 to <27.5 | BMI 27.5 to <30 | BMI 30 to <35 | BMI 35 to <40 | BMI >=40 |
| --- | --- | --- | --- | --- | --- | --- | --- | --- | --- | --- |
| BMI, N= | 457,689 | 2,376 | 8,367 | 46,303 | 94,502 | 109,714 | 84,917 | 80,066 | 22,717 | 8,727 |
| BMI Mean (SD) kg/m2 | 27.42<br>(4.78) | 17.65<br>(0.81) | 19.40<br>(0.42) | 21.48<br>(0.69) | 23.84<br>(0.71) | 26.23<br>(0.71) | 28.67<br>(0.71) | 32.00<br>(1.39) | 36.97<br>(1.38) | 43.76<br>(3.73) |
| WHR, N= | 457,556 | 2,376 | 8,364 | 46,292 | 94,488 | 109,689 | 84,902 | 80,033 | 22,703 | 8,709 |
| WHR Mean (SD) | 0.87<br>(0.09) | 0.77<br>(0.06) | 0.78<br>(0.06) | 0.80<br>(.07) | 0.83<br>(.07) | 0.87<br>(.08) | 0.90<br>(0.08) | 0.92<br>(.09) | 0.93<br>(0.09) | 0.92<br>(0.10) |
| Sex, N= | 457,689 | 2,376 | 8,367 | 46,303 | 94,502 | 109,714 | 84,917 | 80,066 | 22,717 | 8,727 |
| Female | 54.38% | 79.21% | 79.67% | 72.92% | 59.86% | 48.74% | 44.67% | 48.78% | 59.69% | 68.40% |
| Male | 45.62% | 20.79% | 20.33% | 27.08% | 40.14% | 51.26% | 55.33% | 51.22% | 40.31% | 31.60% |
| Days exercise, N= | 433,259 | 2,239 | 7,941 | 44,518 | 90,627 | 104,607 | 80,151 | 74,488 | 20,787 | 7,901 |
| 0 | 37.17% | 44.08% | 36.85% | 32.39% | 31.81% | 34.22% | 37.82% | 43.56% | 51.29% | 59.38% |
| 1 | 14.14% | 13.93% | 14.80% | 15.31% | 15.04% | 14.69% | 13.86% | 12.91% | 11.70% | 10.40% |
| 2 | 15.85% | 12.77% | 15.00% | 16.42% | 17.07% | 16.76% | 15.61% | 14.62% | 12.78% | 10.77% |
| 3 | 13.81% | 9.96% | 12.66% | 14.94% | 15.35% | 14.56% | 13.64% | 12.12% | 10.44% | 8.72% |
| 4 | 6.51% | 4.87% | 6.80% | 7.23% | 7.39% | 6.91% | 6.52% | 5.43% | 4.32% | 3.28% |
| 5 | 6.89% | 6.97% | 6.51% | 7.10% | 7.19% | 7.20% | 7.07% | 6.47% | 5.50% | 4.28% |
| 6 | 2.00% | 1.65% | 2.35% | 2.38% | 2.19% | 2.13% | 1.91% | 1.72% | 1.36% | 1.14% |
| 7 | 3.61% | 5.76% | 5.04% | 4.23% | 3.95% | 3.52% | 3.59% | 3.17% | 2.61% | 2.03% |
| Alcohol frequency N= | 456,669 | 2,363 | 8,335 | 46,205 | 94,344 | 109,485 | 84,746 | 79,848 | 22,643 | 8,700 |
| Daily or almost daily | 20.55% | 22.09% | 22.66% | 22.75% | 22.84% | 22.28% | 20.82% | 17.45% | 12.08% | 7.56% |
| Three or four times a week | 23.07% | 18.74% | 21.82% | 24.29% | 24.81% | 24.81% | 23.71% | 20.78% | 15.97% | 11.41% |
| Once or twice a week | 25.71% | 18.91% | 21.99% | 24.34% | 25.72% | 25.83% | 26.34% | 26.49% | 25.81% | 22.98% |
| One to three times a month | 11.13% | 9.60% | 10.53% | 10.55% | 10.26% | 10.33% | 10.63% | 12.51% | 15.10% | 16.67% |
| Special occasions | 11.54% | 15.32% | 12.42% | 10.39% | 9.57% | 9.89% | 10.81% | 13.68% | 19.30% | 25.10% |
| Never | 8.01% | 15.32% | 10.57% | 7.68% | 6.80% | 6.86% | 7.70% | 9.09% | 11.74% | 16.28% |
| Qualifications, N= | 448,825 | 2,311 | 8,215 | 45,554 | 92,831 | 107,669 | 83,240 | 78,294 | 22,175 | 8,536 |
| College/University degree | 32.31% | 41.32% | 43.29% | 42.01% | 37.66% | 32.99% | 28.90% | 25.32% | 23.13% | 21.88% |
| A levels or equivalent | 11.21% | 13.24% | 13.33% | 12.92% | 11.75% | 11.17% | 10.73% | 10.21% | 10.30% | 10.65% |
| O levels or equivalent | 21.81% | 19.26% | 19.81% | 20.26% | 21.63% | 22.04% | 21.94% | 22.28% | 22.97% | 23.28% |
| CSEs or equivalent | 5.80% | 3.76% | 4.67% | 4.93% | 5.20% | 5.48% | 6.14% | 6.69% | 6.84% | 8.32% |
| NVQ or equivalent | 6.66% | 3.94% | 3.60% | 4.15% | 5.46% | 6.77% | 7.80% | 8.07% | 8.02% | 8.07% |
| Other professional qualifications | 5.21% | 4.24% | 4.25% | 4.68% | 4.84% | 5.17% | 5.46% | 5.72% | 5.64% | 5.37% |
| None of the above | 16.99% | 14.24% | 11.05% | 11.04% | 13.47% | 16.37% | 19.03% | 21.72% | 23.11% | 22.43% |
| Employment, N= | 455,287 | 2,357 | 8,313 | 46,074 | 94,073 | 109,168 | 84,475 | 79,587 | 22,568 | 8,672 |
| In paid employment/self-employed | 57.67% | 52.14% | 59.34% | 61.32% | 59.68% | 57.74% | 56.63% | 55.65% | 54.17% | 52.43% |
| Retired | 33.38% | 29.24% | 27.19% | 28.52% | 32.51% | 34.97% | 35.40% | 34.29% | 32.91% | 29.13% |
| Looking after home/family | 2.80% | 6.50% | 6.09% | 4.61% | 3.12% | 2.27% | 2.09% | 2.29% | 2.78% | 3.54% |
| Unable to work because of ill health | 3.17% | 6.67% | 3.39% | 2.33% | 1.93% | 2.23% | 2.99% | 4.59% | 6.82% | 11.05% |
| Unemployed | 1.69% | 3.31% | 2.11% | 1.52% | 1.40% | 1.54% | 1.76% | 1.93% | 2.24% | 2.69% |
| Doing unpaid work | 0.47% | 1.06% | 0.73% | 0.70% | 0.53% | 0.43% | 0.36% | 0.40% | 0.34% | 0.51% |
| Full or part time student | 0.26% | 0.34% | 0.45% | 0.36% | 0.27% | 0.25% | 0.24% | 0.23% | 0.23% | 0.24% |
| None of the above | 0.57% | 0.64% | 0.71% | 0.64% | 0.57% | 0.56% | 0.54% | 0.62% | 0.50% | 0.42% |
| Quintile of deprivation, N= | 457,156 | 2,371 | 8,358 | 46,251 | 94,386 | 109,606 | 84,815 | 79,968 | 22,689 | 8,712 |
| 1 (lowest deprivation) | 20.06% | 17.63% | 18.95% | 20.95% | 21.90% | 21.56% | 20.03% | 17.98% | 14.59% | 11.81% |
| 2 | 20.00% | 17.17% | 18.09% | 19.92% | 21.21% | 20.69% | 20.51% | 18.92% | 16.93% | 13.88% |
| 3 | 20.0% | 16.91% | 19.19% | 20.06% | 20.16% | 20.58% | 20.24% | 19.67% | 18.37% | 17.57% |
| 4 | 20.01% | 21.00% | 20.81% | 20.12% | 19.41% | 19.58% | 19.89% | 20.51% | 21.80% | 21.91% |
| 5 (highest deprivation) | 19.93% | 27.29% | 22.96% | 18.95% | 17.32% | 17.59% | 19.33% | 22.91% | 28.31% | 34.83% |
| Age at recruitment, N= | 457,689 | 2,376 | 8,367 | 46,303 | 94,502 | 109,714 | 84,917 | 80,066 | 22,717 | 8,727 |
| Age (years) at recruitment Mean (SD) | 56.52<br>(8.10) | 55.40<br>(8.15) | 54.53<br>(8.3) | 54.98<br>(8.27) | 56.14<br>(8.17) | 56.89<br>(8.08) | 57.13<br>(8.03) | 57.05<br>(7.91) | 56.58<br>(7.82) | 55.44<br>(7.71) |

Notes to table: SD: Standard deviation. BMI: Body mass index. WHR: Waist hip ratio NVQ: National Vocational Qualification. CSE: Certificate of Secondary Education

Some 54.4% of individuals included in the analysis sample had positive healthcare costs over the period of follow-up. Mean BMI overall was 27.4 kg/m2 (SD: 4.8); BMI amongst men was 27.8 kg/m2 (SD: 4.2) and amongst women 27.1 kg/m2 (SD 5.2). The proportion of participants with BMI >30kg/m2 was 24.4%, with more men (25.3%) than women (23.5%) meeting the definition of obesity. This is slightly lower than the prevalence of obesity in England in 2010 (the last year of recruitment to the study) amongst all adults of 26.1% (26.2% males, 26.1% females) (36).

There was limited missingness (<0.1%) amongst baseline covariates used in the adjusted analysis, with the exception of the exercise variable (5.4% coded as missing) and the qualifications variable (1.9% missing). Tests of family and link functions for the GLM supported the use of a Gamma family with logarithmic link when modelling the cost outcome. Further details on the results of these tests are provided in supplementary material.

All models found a positive association between measures of adiposity and hospital costs (Table 2). Marginal effects represent a specific change in the outcome associated with each scenario, leaving other variables at their observed levels, and averaged over all individuals in the sample. For example, an increase in categorical BMI from the base category of 25 kg/m2 to 27.5kg/m2 to the 30 kg/m2 to 35kgm2 category is associated with an increase in costs of £106.24 (99% confidence interval: £91.59 to £120.90). This marginal effect can be interpreted as the hypothetical difference if all individuals differed only in their BMI category, but not on any other covariate. These average adjusted predictions reflect the expected values of the inpatient cost outcome for specific sets of values for the predictor variables, such as., for example, the effect of a standard deviation increase in BMI.

**Table 2.** Marginal effects and average adjusted predicted inpatient hospital costs.

|  | Marginal effect | 99% confidence interval | Average adjusted predicted cost | 99% confidence interval |
| --- | --- | --- | --- | --- |
| Outcome: Continuous BMI |  |  |  |  |
| Effect of unit change in BMI | £13.61 | £12.60 to £14.63 | £468.30 | £464.84 to £471.77 |
| Effect of SD change in BMI | £69.20 | £64.98 to £73.42 | £537.51 | £531.69 to £543.32 |
| Outcome: Binary BMI |  |  |  |  |
| Effect of change in BMI-defined obesity status | £136.35 | £124.62 to £148.08 | £569.25 | £561.51 to £576.99 |
| Outcome: Categorical BMI |  |  |  |  |
| 10 to <18.5 kg/m<sup>2</sup> | £41.62 | -£77.86 to £161.10 | £473.35 | £354.31 to £592.39 |
| 18.5 to <20 kg/m<sup>2</sup> | -£31.94 | -£68.91 to £ 5.03 | £399.79 | £364.04 to £435.55 |
| 20 to <22.5 kg/m<sup>2</sup> | -£40.94 | -£57.02 to -£24.86 | £390.79 | £377.44 to £404.15 |
| 22.5 to <25 kg/m<sup>2</sup> | -£20.83 | -£34.52 to -£7.15 | £410.90 | £400.65 to £421.15 |
| 25 to <27.5 kg/m<sup>2</sup> (Base category) | - | - | £431.73 | £422.91 to £440.56 |
| 27.5 to <30 kg/m<sup>2</sup> | £46.01 | £31.39 to £60.63 | £477.74 | £466.05 to £489.43 |
| 30 to <35 kg/m<sup>2</sup> | £106.24 | £91.59 to £120.90 | £537.98 | £526.52 to £549.44 |
| 35 to <40 kg/m<sup>2</sup> | £188.99 | £162.37 to £215.61 | £620.72 | £595.83 to £645.62 |
| >= 40 kg/m<sup>2</sup> | £300.14 | £257.60 to £342.69 | £731.87 | £690.43 to £773.32 |
| Outcome: Continuous WHR |  |  |  |  |
| Effect of 0.01 change in WHR | £ 8.79 | £ 8.20 to £ 9.38 | £476.51 | £472.91 to £480.12 |
| Effect of SD change in WHR | £85.06 | £78.93 to £91.18 | £552.77 | £545.25 to £560.30 |
| Outcome: Binary WHR |  |  |  |  |
| Effect of change in regional adiposity status | £96.78 | £85.78 to £105.87 | £514.16 | £508.50 to £519.82 |

Notes to table: SD: Standard deviation. BMI: Body mass index. WHR: Waist hip ratio

Marginal effects for continuous outcomes require the specification of a particular “margin” of change, but otherwise have a similar interpretation of the marginal effects of categorical outcomes. For example, the effect of a one standard deviation increase in BMI is to increase in inpatient hospital costs by £69.20 (99% CI: £64.98 to £73.42).

The relative effect of a unit change in continuous BMI amounts to approximately 2.9% (99% CI: 2.7% to 3.2%) in increased costs per person year of follow-up. Relative effects for the categorial outcomes are presented in supplementary material.

Figure 2 displays average adjusted predicted costs at representative values of continuous BMI, in essence capturing predictions at unit increments of BMI from the small to the large BMI integer values. Figure 3 presents the same outcome according to age, and stratified by sex to examine whether these variables may influence inpatient costs. The point estimates and confidence intervals are similar, and no difference in sex is apparent at any age. However, there is a clear gradient with respect to age, with older members of the cohort predicted to have higher inpatient costs. Figure 4 summarises average adjusted predictions for the categorical BMI outcome. There is some evidence of a J-shaped relationship in point estimates for lower levels of categorical BMI, although this evidence is weak given the widths of the confidence intervals around these estimates.

**Figure 2.**
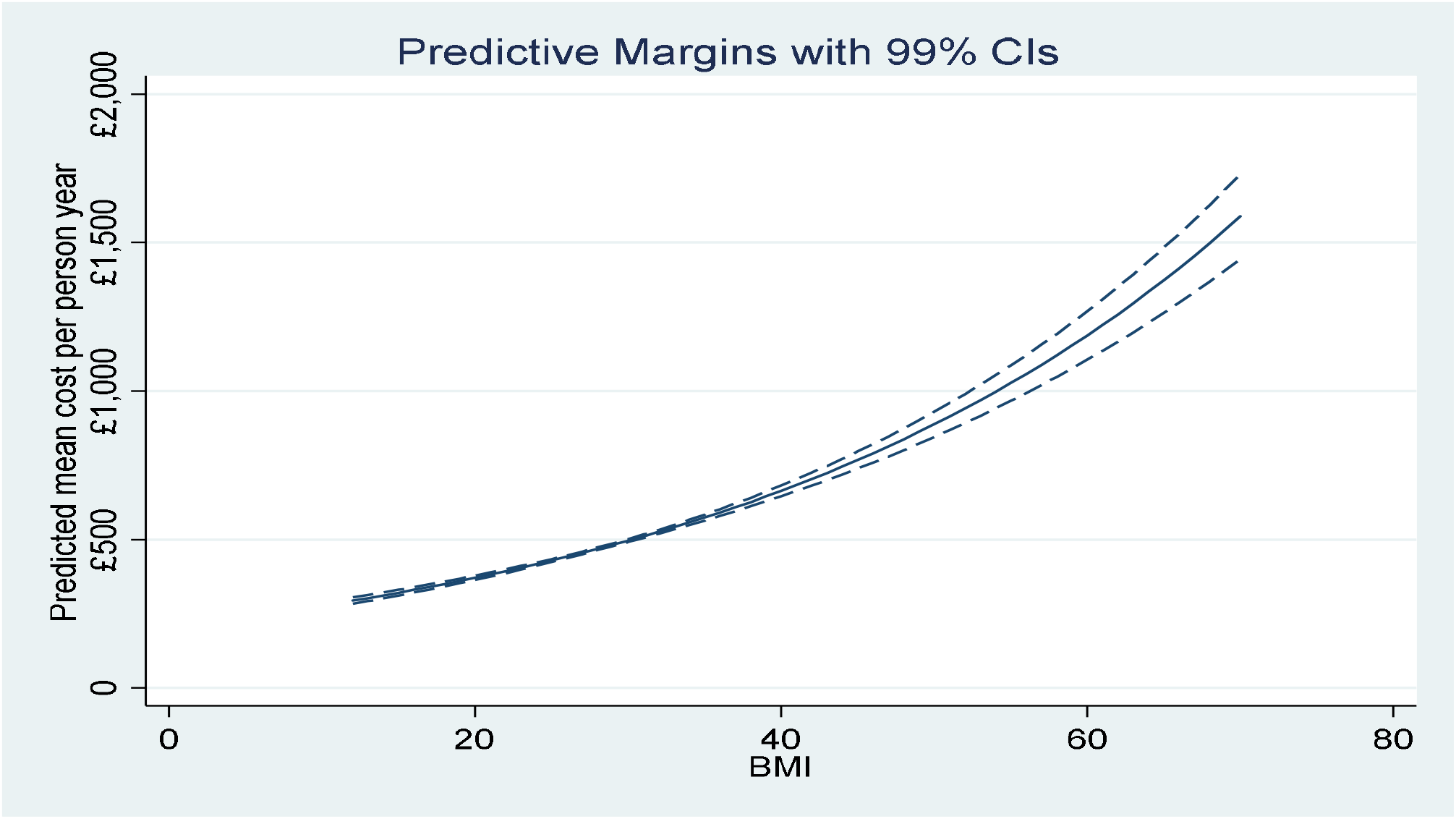
Average adjusted predicted healthcare costs by continuous BMI.

**Figure 3.**
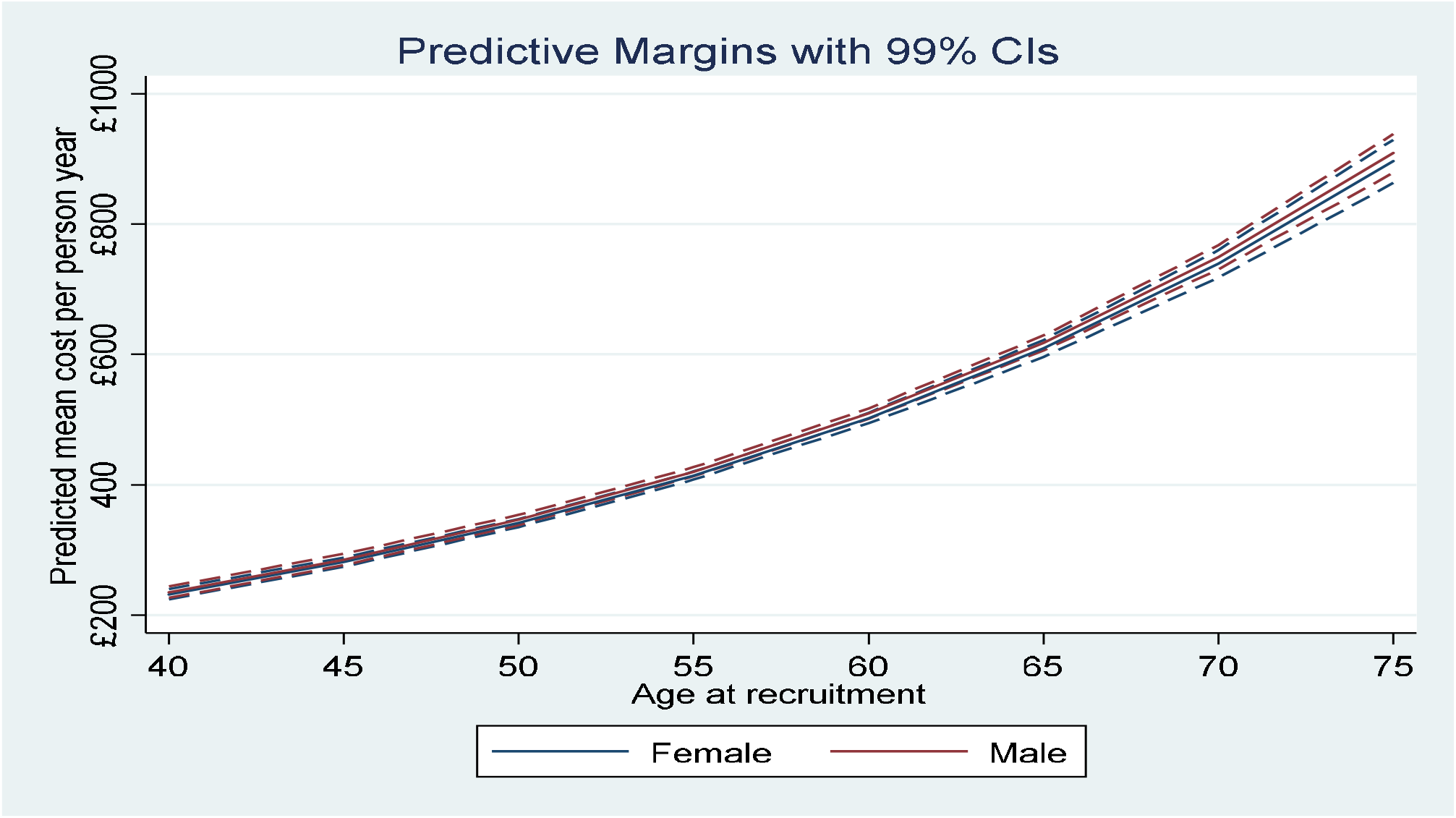
Average adjusted predicted healthcare costs by continuous BMI, by age and stratified by sex.

**Figure 4.**
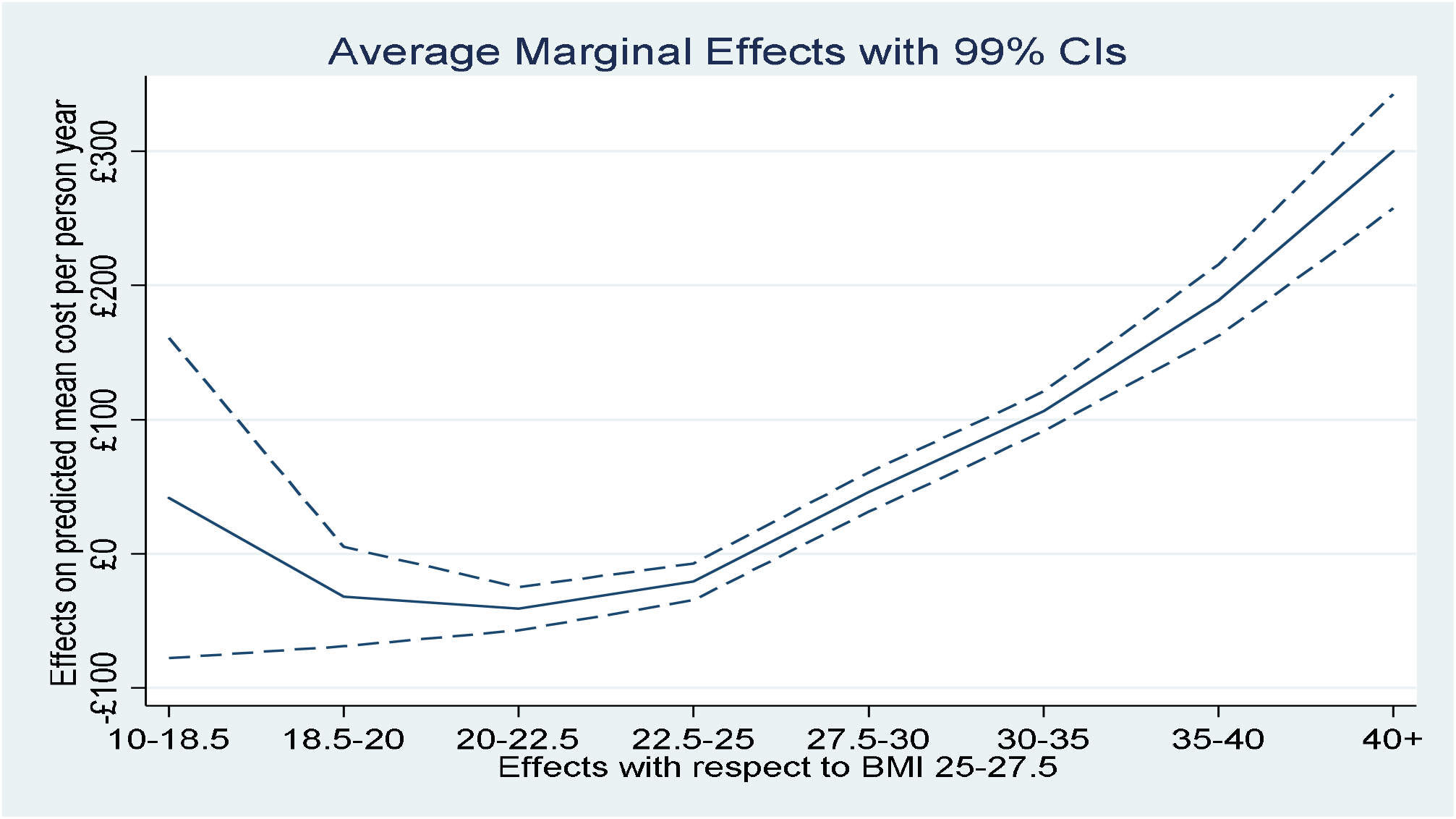
Average marginal effects by categorical BMI.

Three of the five specifications assessed in sensitivity analysis were very similar to the base case predictions (Figure 5). The effect of BMI attenuates slightly when controlling for waist hip ratio. Including only never smokers slightly reduced the estimated marginal effect to £13.10 (99% confidence interval: £11.77 to £14.43), and predicted costs are similar to the base model near mean BMI. Excluding individuals who reported no pre-existing condition health conditions had the largest impact of all scenarios assessed, reducing the marginal effect of a unit change of BMI to £5.51 (99% confidence interval: £4.13 to £6.89). Supplementary material describes average adjusted predictions over the range of BMI for these sensitivity analyses Results from sensitivity analysis of the other outcomes reported in Table 2 were similar.

**Figure 5.**
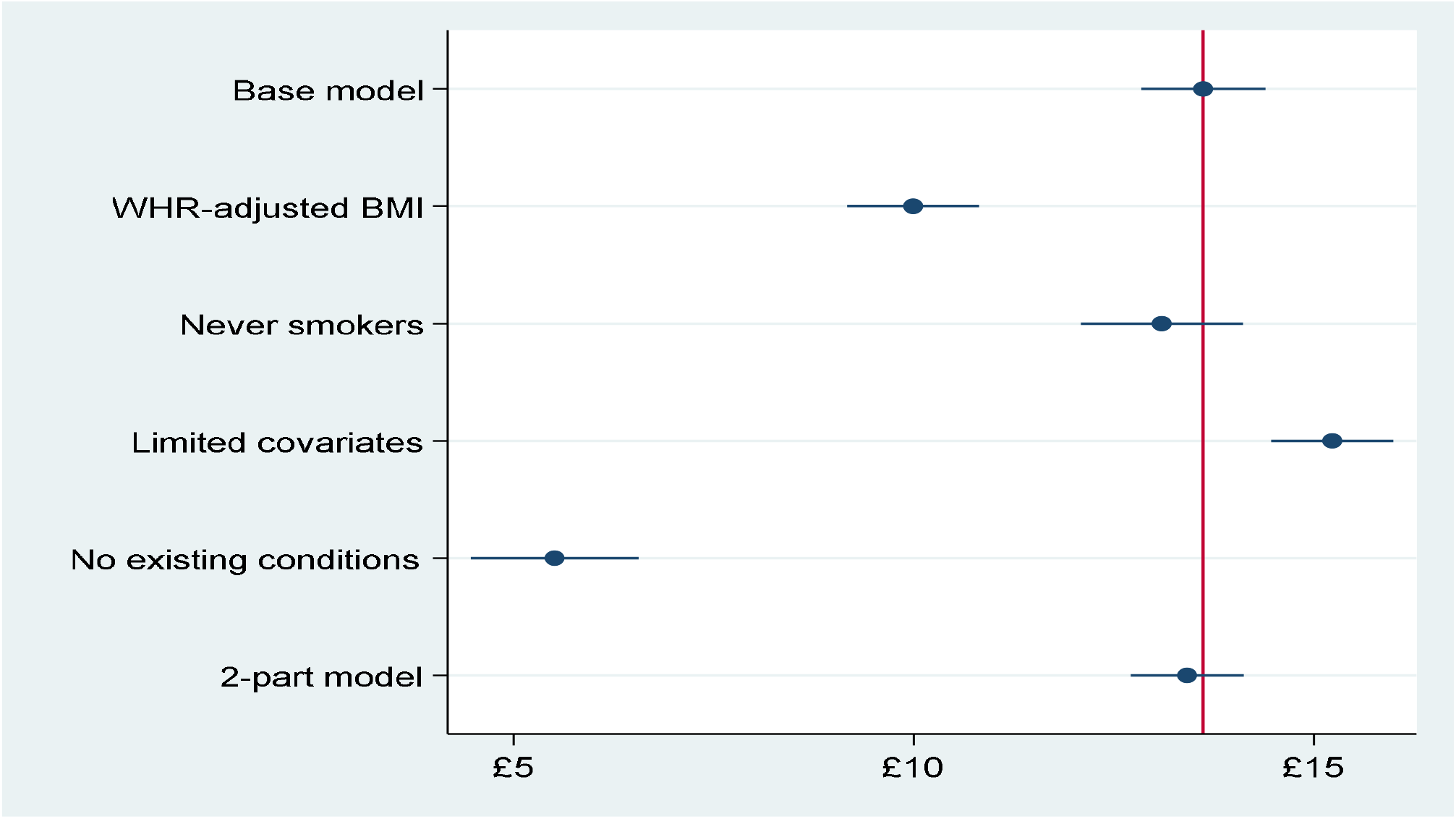
Average marginal effects per unit change in continuous BMI.

### Discussion

Higher rates of adiposity are associated with higher inpatient hospital costs in this cohort of middle-aged and older individuals. This is broadly consistent with findings from studies with similar designs. A systematic review (37) of the association between total annual healthcare costs and BMI in studies using individual participant data found progressive increases in annual hospital costs for increasing levels of obesity, a finding demonstrated here in all models and for all outcomes. The systematic review also reported that overweight and obesity increased median inpatient hospital costs relative to costs associated with BMI of 18.5kg/m2 to <25kgm2 by 12% and 34% respectively. These figures are similar to those reported here, for which the comparable relative increases in cost for BMI 25kg/m2 to <27.5 kg/m2 relative to 22.5 to 25kg/m2 of 16%, and 31% for BMI >=30kg/m2 relative to 22.5 to <25kg/m2.

Korda et al studied hospital costs amongst 224,254 Australian adults aged at least 45 years, and found an increase in costs of between 14% and 24% (depending on category of age) for BMI between 27.5-30 kg/m2 relative to 22.5-<25kg/m2, rising to 77% to 115% for BMI between 40-50 kg/m2. The comparable increase in costs for individuals of all ages in UK Biobank >=40kgm2 is 78%. The findings are also broadly similar to the US study of Andreyeva et al (38).

Marginal effects for the continuous BMI outcome are similar to the observational estimates in Cawley and Meyerhoefer for US adults of $49 in 2005 US dollars per unit change in BMI, although their estimate of the marginal effect of obesity ($656) is much higher than the effect reported in Table 2 for the UK Biobank cohort. Predicted costs and marginal effects are estimated to be somewhat lower than in the large prospective women-only study of Kent et al (20), perhaps reflecting the relatively “healthier and wealthier” composition of participants in UK Biobank.

### Strengths and limitations

This analysis benefitted from access to a large volume of patient level data obtained from a prospective cohort study with comprehensive information on a variety of baseline characteristics. Measures of BMI and related variables were obtained by research staff, and the errors and biases associated with self-report of weight and height were avoided (39). The use of traditional and impedance-based measures of BMI offered a degree of further validation for the core exposure. The analysis included both continuous and categorical/binary outcomes of weight-related measures, in contrast to much of the literature on these associations which is generally restricted to categorical or binary classifications of BMI.

The analysis has a number of limitations. The analysis presented in this paper is observational and cannot reveal the causal association between measures of adiposity and inpatient costs. For example, the association between costs and BMI may be confounded by the effect of unmeasured baseline health conditions that affect both BMI and hospital costs. Adjustment for baseline pre-existing conditions in sensitivity analysis revealed a relatively large moderating effect, but only captures known health conditions that were accurately reported at baseline.

The results could therefore be affected by some degree of reverse causality, the precise degree of which cannot be uncovered in this type of observational analysis. The results from excluding individuals without pre-existing conditions was suggestive – effect sizes attenuated but did not encompass the null. Nevertheless, this is necessarily a partial and incomplete test of reverse causality.

A further channel for costs to influence BMI may be treatment regimens that affect weight (such as selective serotonin reuptake inhibitors for depression), or treatment decisions that used BMI thresholds (such as for bariatric surgery) as a basis for intervention. However, observing these associations may give the appearance of the direction of causality running from healthcare costs to BMI, but in reality are better conceived of as further instances of confounding (by depression in the first example) rather than an actual causal effect on the level of BMI from the level of costs per se.

Methods for causal inference, such as Mendelian Randomization – the use of germline genetic variants as instrument variables for exposures such as BMI – or the use of BMI of a biological relative as an instrument for own BMI, offer promise in this area as a means to overcome confounding (40). Complementary study designs and triangulation (21) between these different designs will almost certainly be necessary even after causal analyses of the adiposity/cost relationship.

Biobank cohort participants differ from the general population: they are more likely to be older, female and university graduates. They are less likely to be deprived, obese, to smoke, to drink alcohol on a daily-basis, or to have self-reported health conditions. There is evidence of a selection bias from a “healthy volunteer” effect (22). The rate of all-cause mortality is 46% lower in the cohort than in the population from which it was drawn (23). This may induce associations between outcomes and participant characteristics when none exist in truth due to the operation of collider bias caused by the self-selection of a healthier and more educated population into the cohort (41).

This is likely to impair the representativeness and generalisability of the findings of this study to the wider UK population from which the cohort is drawn. The direction of the bias that this imparts is unknown. However, it is plausible that the direction of bias may be downwards, suggesting that the estimates presented here are likely to be an underestimate of the effect of BMI on healthcare costs. This would occur if, for example, the healthy behaviours and favourable circumstances of cohort participants (relative to the general population) are related to unmeasured variables that themselves tend to mitigate the consequences of higher BMI. Robust causal methods are required to resolve this uncertainty.

Modelling cost outcomes is challenging, not least because methods to identify appropriate link and family functions in GLM have limitations and require a degree of contextual interpretation, and mis-specification is possible despite robust standard errors being used for all models. The costs assigned to each individual episode of care are based on HRGs that are essentially averages of costs of types of care, and may conceal individual variation in actual resources used.

Analysis did not encompass all healthcare costs. Primary care data is not linked to UK Biobank, and hospital data is restricted to inpatient care, although Kent et al (20) report that 30-50% of total overweight and obesity attributable costs relate to inpatient care. It is also probable that many costs will be correlated with the same direction of effect across different categories of provider care, albeit that the scale of effects is likely to differ, This question requires linked data that is not available for the UK Biobank cohort. The remuneration of Scottish hospitals differs from that in England and Wales, and Scottish hospitalisation data could not be combined with that from the latter two jurisdictions.

The dataset includes private patients treated in NHS hospitals, but not private patients treated elsewhere. Approximately 37% of the cohort answered a question on use of private health insurance, of whom 2.4% self-report exclusive use of private healthcare. Nevertheless, 39% of individuals reporting exclusive use of private healthcare have non-zero NHS inpatient costs. Further data on reported use of private healthcare in the cohort is presented in supplementary material.

## Conclusion

BMI is a modifiable risk factor for a variety of healthcare conditions. Evidence from analysis of the UK Biobank is consistent with findings from other cohorts in demonstrating a robust association between BMI and inpatient hospital costs. However, evidence from randomised study designs or causal inference on observational data is required to validate these findings and to inform public policy toward excess adiposity.

## Acknowledgement

This research has been conducted using the UK Biobank Resource under Application Number 29294.

